## Supplementary information for "FLASH-MM: fast and scalable single-cell differential expression analysis using linear mixed-effects models"

**References** **16**

### 1 Linear mixed-effects models

In this section, we first describe the linear mixed-effects model (LMM) and then introduce the methods for the LMM parameter estimation, hypothesis testing, prediction of random effects, and gradient algorithms for computing the LMM variance components, see Searle, Casella, and McCulloch (2006), Jiang (2007) and references therein for details.

A linear mixed-effects model (LMM) is an extension of general linear model, which contains both fixed effects and random effects as expressed below (Searle, Casella, and McCulloch 2006)

$$y = X\beta + Zb + \epsilon, \tag{1}$$

where $y$ is an $n \times 1$ vector of observations, $X$ is an $n \times p$ design matrix for fixed effects $\beta$, $Z$ is an $n \times q$ design matrix for random effects $b$, and $\epsilon$ is an $n \times 1$ vector of residual errors. The term of random effects may be a combination of various random-effect components:

$$Zb = Z_1 b_1 + \cdots + Z_K b_K,$$

where $Z = [Z_1, \ldots, Z_K]$, $b = [b_1^T, \ldots, b_K^T]^T$, $K$ is the number of random-effect components, $Z_k$ is an $n \times q_k$ design matrix for the $k$-th random-effect component, and $\sum_{k=1}^{K} q_k = q$. The superscript $T$ denotes a transpose of vector or matrix. The basic assumptions are as follows:

(1) The design matrix $X$ is of full rank, satisfying conditions of estimability for the parameters.
(2) The random vectors $b_k$ and $\epsilon$ are independent and follow a normal distribution, $b_k \sim N(\mathbf{0}, \sigma_k^2 I_{q_k})$ and $\epsilon \sim N(\mathbf{0}, \sigma^2 I_n)$, where $\sigma_k^2$ and $\sigma^2$ are unknown parameters, called variance components, $\mathbf{0}$ is a vector or matrix of zero elements, and $I_n$ is an $n \times n$ identity matrix.

Assumption (1) implies $p < n$. We also assume $q_k < n$. If $q_k \geq n$, we can use principal component analysis (PCA) to obtain an equivalent LMM with the number of random effects less than $n$, as described late by (47).

Hartley and Rao (1967) specified the LMM (1) as mixed analysis of variance (ANOVA) models. Harville (1977) introduced the general linear mixed-effects models with covariance matrices $Cov(b) = D_\phi$ and $Cov(\epsilon) = D_\psi$, specified by the unobservable parameter vectors $\phi$ and $\psi$. Laird and Ware (1982) described the linear mixed-effects models for longitudinal data, repeated measures data, or grouped data.

The random effects reflect variations between groups (subjects) and correlations within groups (subjects). Suppose $Cov(b) = \sigma_b^2 I_q$. Then the variance of the $i$th observation and the correlation between two observations $i$ and $j$ are

$$Var(y_i) = \sigma^2 + \sigma_b^2 z_i^T z_i,$$
$$Cor(y_i, y_j) = \frac{\sigma_b^2 z_i^T z_j}{(\sigma^2 + \sigma_b^2 z_i^T z_i)^{1/2}(\sigma^2 + \sigma_b^2 z_j^T z_j)^{1/2}},$$

where $z_i$ is the $i$th row of $Z$ corresponding to the $i$th observation. If two observations $i$ and $j$ come from different subjects and $z_i^T z_i \neq z_j^T z_j$, then $Var(y_i) \neq Var(y_j)$, reflecting the variation between subjects (inter-subject variability). If the two observations come from same subject, usually $z_i = z_j$ and then $Cor(y_i, y_j) > 0$, that is, the two observations within a subject are correlated, reflecting the intra-subject correlation.

#### 1.1 LMM parameter estimation

Hartley and Rao (1967) developed maximum likelihood (ML) method for estimating the LMM parameters, that is, the fixed effects and variance components. The ML method estimates all parameters of fixed effects and variance components together. Patterson and Thompson (1971) proposed a modified maximum likelihood procedure which partitions the data into two mutually uncorrelated parts, one being free of the fixed effects used for estimating variance components, called restricted maximum likelihood (REML) estimators. The REML estimator is unbiased, and the ML estimator (MLE) of variance components is biased in general. For given variance components, both ML and REML methods provide with the same estimates of fixed effects. In this section, we briefly describe both ML and REML methods, see Hartley and Rao (1967), Patterson and Thompson (1971), Harville (1977), and Jiang (2007) for the details.

##### 1.1.1 Maximum likelihood estimation

Under the assumptions that random vectors $b_i$ and $\epsilon$ are independent and have a normal distribution, $b_i \sim N(\mathbf{0}, \sigma_i^2 I_{q_i})$ and $\epsilon \sim N(\mathbf{0}, \sigma^2 I_n)$, we have $y \sim N(X\beta, V_\theta)$, with a probability density function (pdf)

$$f(y|\beta, \theta) = \frac{1}{\sqrt{(2\pi)^n \det(V_\theta)}} \exp[-\frac{1}{2}(y - X\beta)^T V_\theta^{-1}(y - X\beta)], \tag{2}$$

where

$$\theta = [\theta_0, \theta_1, \ldots, \theta_K]^T = [\sigma^2, \sigma_1^2, \ldots, \sigma_K^2]^T,$$
$$V_\theta = \sigma^2 I_n + \sigma_1^2 Z_1 Z_1^T + \ldots + \sigma_K^2 Z_K Z_K^T = \sigma^2 I_n + Z D_\theta Z^T,$$
$$D_\theta = \begin{bmatrix} \theta_1 I_{q_1} & \ldots & \mathbf{0} \\ \vdots & \ddots & \vdots \\ \mathbf{0} & \ldots & \theta_K I_{q_K} \end{bmatrix}.$$

The maximum likelihood estimation (MLE) is obtained by maximizing the log-likelihood

$$l(\beta, \theta) = \log f(y|\beta, \theta) = -\frac{1}{2} n \log 2\pi - \frac{1}{2} \log\det(V_\theta) - \frac{1}{2}(y - X\beta)^T V_\theta^{-1}(y - X\beta). \tag{3}$$

The first derivatives of the log-likelihood are

$$\begin{cases} \frac{\partial l}{\partial \beta} = X^T V_\theta^{-1} y - X^T V_\theta^{-1} X\beta, \\ \frac{\partial l}{\partial \theta_i} = -\frac{1}{2} tr(V_\theta^{-1} V_i) + \frac{1}{2}(y - X\beta)^T V_\theta^{-1} V_i V_\theta^{-1}(y - X\beta), \ i = 0, \ldots, K, \end{cases} \tag{4}$$

where $V_i = Z_i Z_i^T$ and $Z_0 = I_n$. By equating the first derivatives to zero, we have the MLE equations:

$$\begin{cases} X^T V_\theta^{-1} X\beta = X^T V_\theta^{-1} y, \\ tr(V_\theta^{-1} V_i) = y^T R_\theta V_i R_\theta y, \ i = 0, \dots, K, \end{cases} \tag{5}$$

where

$$R_\theta = V_\theta^{-1} - V_\theta^{-1} X (X^T V_\theta^{-1} X)^{-1} X^T V_\theta^{-1}. \tag{6}$$

With $\theta$ given, from (5), we have the MLE of $\beta$ and the covariance matrix:

$$\hat{\beta} = (X^T V_\theta^{-1} X)^{-1} X^T V_\theta^{-1} y, \tag{7}$$

$$var(\hat{\beta}) = (X^T V_\theta^{-1} X)^{-1}.$$

Note

$$P_{V_\theta^{-1/2} X} = V_\theta^{-1/2} X (X^T V_\theta^{-1} X)^{-1} X^T V_\theta^{-1/2}$$

is a projection matrix, and $R_\theta = V_\theta^{-1/2}(I_n - P_{V_\theta^{-1/2}X}) V_\theta^{-1/2}$ is a residual (maker) matrix. The following properties can be readily verified:

$$R_\theta X = \mathbf{0}, \quad R_\theta V_\theta R_\theta = R_\theta. \tag{8}$$

The information matrix is given by the second derivatives of $l(\beta, \theta)$

$$\begin{cases} \frac{\partial^2 l}{\partial \beta \partial \beta^T} = -X^T V_\theta^{-1} X \\ \frac{\partial^2 l}{\partial \beta \partial \theta_i} = -X^T V_\theta^{-1} V_i V_\theta^{-1} (y - X\beta) \\ \frac{\partial^2 l}{\partial \theta_i \partial \theta_j} = \frac{1}{2} tr(V_\theta^{-1} V_i V_\theta^{-1} V_j) - (y - X\beta)^T V_\theta^{-1} V_i V_\theta^{-1} V_j V_\theta^{-1} (y - X\beta) \end{cases} \tag{9}$$

Using $E(y - X\beta) = \mathbf{0}$ and $E[(y - X\beta)^T A (y - X\beta)] = tr(A V_\theta)$ for any $n \times n$ matrix $A$, we have Fisher information matrix:

$$I(\beta, \theta) = -\begin{bmatrix} E(\frac{\partial^2 l}{\partial \beta \partial \beta^T}) & E(\frac{\partial^2 l}{\partial \beta \partial \theta^T}) \\ E(\frac{\partial^2 l}{\partial \theta \partial \beta^T}) & E(\frac{\partial^2 l}{\partial \theta \partial \theta^T}) \end{bmatrix} = \begin{bmatrix} var(\hat{\beta})^{-1} & \mathbf{0} \\ \mathbf{0} & I(\theta) \end{bmatrix}, \tag{10}$$

where

$$var(\hat{\beta}) = (X^T V_\theta^{-1} X)^{-1},$$

$$I(\theta) = -E\Big(\frac{\partial^2 l}{\partial \theta \partial \theta^T}\Big) = \big\{\frac{1}{2} tr(V_\theta^{-1} V_i V_\theta^{-1} V_j)\big\}_{0 \le i,j \le K}.$$

Recall that $I_n$ denotes an $n \times n$ identity matrix. Without confusion, $I(\cdot)$ denotes a function of the Fisher information matrix with corresponding parameters as variables. The Fisher information matrix may also be written as

$$I(\beta, \theta) = E\left[\begin{pmatrix} \frac{\partial l}{\partial \beta} \\ \frac{\partial l}{\partial \theta} \end{pmatrix} \begin{pmatrix} \frac{\partial l}{\partial \beta} \\ \frac{\partial l}{\partial \theta} \end{pmatrix}^T\right],$$

which is a positive semidefinite matrix.

Let $\gamma_i = \sigma_i^2 / \sigma^2$, $i = 1, \dots, K$. Then $V_\theta = \sigma^2 V_\gamma$, where

$$V_\gamma = I_n + \gamma_1 Z_1 Z_1^T + \ldots + \gamma_K Z_K Z_K^T = I_n + Z D_\gamma Z^T. \tag{11}$$

From Weinstein–Aronszajn identity:

$$\det(I_n + AB^T) = \det(I_q + B^T A) \tag{12}$$

for any $n \times q$ matrices $A$ and $B$, we have

$$\det(V_\theta) = \sigma^{2n} \det(I_n + Z D_\gamma Z^T) = \sigma^{2n} \det(I_q + D_\gamma Z^T Z).$$

Note $V_\theta^{-1}(y - X\hat{\beta}) = R_\theta y$. Given the MLEs, $\hat{\beta}$ and $\hat{\theta}$, the maximum log-likelihood (3) can be computed by

$$l(\hat{\beta}, \hat{\theta}) = -\frac{n}{2}\log(2\pi\hat{\sigma}^2) - \frac{1}{2}\log\det(I_q + D_{\hat{\gamma}}Z^TZ) - \frac{1}{2}y^TR_{\hat{\theta}}V_{\hat{\theta}}R_{\hat{\theta}}y, \tag{13}$$

where $\hat{\gamma}_i = \hat{\sigma}_i^2/\hat{\sigma}^2$. From the MLE equations (5),

$$y^TR_{\hat{\theta}}V_{\hat{\theta}}R_{\hat{\theta}}y = \sum_{i=0}^{K}\hat{\sigma}_i^2 y^TR_{\hat{\theta}}V_iR_{\hat{\theta}}y = \sum_{i=0}^{K}\hat{\sigma}_i^2 tr(V_{\hat{\theta}}^{-1}V_i) = n.$$

Then the maximum log-likelihood (13) is reduced as

$$l(\hat{\beta}, \hat{\theta}) = -\frac{n}{2} - \frac{n}{2}\log(2\pi\hat{\sigma}^2) - \frac{1}{2}\log\det(I_q + D_\gamma Z^TZ). \tag{14}$$

From $y^TR_\theta V_\theta R_\theta y = \sigma^{-2}y^TR_\gamma V_\gamma R_\gamma y$ and $R_\gamma y = V_\gamma^{-1}(y - X\hat{\beta})$, we also have

$$\hat{\sigma}^2 = \frac{1}{n}y^TR_{\hat{\gamma}}V_{\hat{\gamma}}R_{\hat{\gamma}}y = \frac{1}{n}(y - X\hat{\beta})^TV_{\hat{\gamma}}^{-1}(y - X\hat{\beta}). \tag{15}$$

**Remark**: The Weinstein–Aronszajn identity (12) can be derived from the equality

$$\begin{bmatrix} I_n & \mathbf{0} \\ B^T & I_q \end{bmatrix} \begin{bmatrix} I_n + AB^T & A \\ \mathbf{0} & I_q \end{bmatrix} \begin{bmatrix} I_n & \mathbf{0} \\ -B^T & I_q \end{bmatrix} = \begin{bmatrix} I_n & A \\ \mathbf{0} & I_q + B^TA \end{bmatrix}.$$

##### 1.1.2 Restricted maximum likelihood

Let $Q$ be an $n \times (n-p)$ full column rank matrix such that $Q^TX = \mathbf{0}$, and $L = V_\theta^{-1}X$, an $n \times p$ full column rank matrix. Then the data y can be partitioned into two parts: $z = Q^Ty$ and $u = L^Ty$. The data $z$ and $u$ are uncorrelated since $Cov(z, u) = Q^TV_\theta L = Q^TX = \mathbf{0}$. From (1),

$$z = Q^TZb + Q^T\epsilon \sim N(\mathbf{0}, Q^TV_\theta Q).$$

The transformed data, $z$, does not contain the fixed effects $\beta$. Based on the log-likelihood function of $z$,

$$l_R(\theta) = -\frac{n-p}{2}\log 2\pi - \frac{1}{2}\log\det(Q^TV_\theta Q) - \frac{1}{2}y^TQ(Q^TV_\theta Q)^{-1}Q^Ty,$$

similar to the MLE equations (5), we have the restricted maximum likelihood (REML) equations:

$$tr[(Q^TV_\theta Q)^{-1}Q^TV_iQ] = y^TQ(Q^TV_\theta Q)^{-1}Q^TV_iQ(Q^TV_\theta Q)^{-1}Q^Ty,\ i = 0, \ldots, K. \tag{16}$$

The REML equations can be further simplified. Let

$$P_A = A(A^TA)^{-1}A^T$$

be the projection matrix for a full column rank matrix $A$. Based on $Q^TX = \mathbf{0}$ and both $X$ and $Q$ are of full rank, for an $n \times n$ positive definite matrix $V$, we have the following equation (Jiang 2007)

$$P_{V^{1/2}Q} = I_n - P_{V^{-1/2}X},$$

that is,

$$Q(Q^TVQ)^{-1}Q^T = V^{-1} - V^{-1}X(X^TV^{-1}X)^{-1}X^TV^{-1}. \tag{17}$$

With (17), the REML equations (16) reduces to

$$tr(R_\theta V_i) = y^TR_\theta V_iR_\theta y,\ i = 0, \ldots, K, \tag{18}$$

where $R_\theta$ is defined as (6),

$$R_\theta = V_\theta^{-1} - V_\theta^{-1}X(X^TV_\theta^{-1}X)^{-1}X^TV_\theta^{-1}.$$

The REML equations (18) do not contain $Q$. Thus the REML estimator does not depend on $Q$.

Let $Q = (I_n - P_X)C$, where $C$ is an $n \times (n-p)$ matrix. Thus

$$z = Q^T y = C^T (I_n - P_X) y = C^T (y - X\hat{\beta}_{LS}),$$

where $\hat{\beta}_{LS} = (X^T X)^{-1} X^T y$ is the least square estimator of $\beta$, is a linear combination of residuals obtained after fitting the fixed effects. So the $z$ is also called error contrasts (Harville 1977).

The REML equations do not include the fixed effects. The fixed effects can be estimated based on the second part of the data: $u = L^T y$, where $L = V_\theta^{-1} X$. From (1),

$$u = L^T X\beta + L^T Z b + L^T \epsilon \sim N(L^T X\beta, L^T V_\theta L).$$

With $\theta$ fixed, based on the data $u$, the MLE of $\beta$ is given by

$$\hat{\beta} = [X^T L (L^T V_\theta L)^{-1} L^T X]^{-1} [X^T L (L^T V_\theta L)^{-1} L^T y] = (X^T V_\theta^{-1} X)^{-1} X^T V_\theta^{-1} y. \tag{19}$$

For a given $\theta$, the MLE of $\beta$ based on $u$ is exactly the same with that based on $y$ in (7).

The first and second derivatives of $l_R$ are given as follows:

$$\frac{\partial l_R}{\partial \theta_i} = -\frac{1}{2} tr(R_\theta V_i) + \frac{1}{2} y^T R_\theta V_i R_\theta y, \tag{20}$$

$$\frac{\partial^2 l_R}{\partial \theta_i \partial \theta_j} = \frac{1}{2} tr(R_\theta V_i R_\theta V_j) - y^T R_\theta V_i R_\theta V_j R_\theta y. \tag{21}$$

From (21), using $E(y^T A y) = tr(A V_\theta) + \beta^T X^T A X \beta$, $R_\theta X = \mathbf{0}$ and $R_\theta V_\theta R_\theta = R_\theta$, we have Fisher information matrix:

$$I(\theta) = -E\Big(\frac{\partial^2 l_R}{\partial \theta \partial \theta^T}\Big) = \big\{\frac{1}{2} tr(R_\theta V_i R_\theta V_j)\big\}_{0 \le i,j \le K}. \tag{22}$$

Recall $V_\theta = \sigma^2 (I_n + Z D_\gamma Z^T)$. From (17) and the Weinstein–Aronszajn identity (12),

$$\begin{aligned} \det(Q^T V_\theta Q) &= \det[\sigma^2 (Q^T Q + Q^T Z D_\gamma Z^T Q)] \\ &= \sigma^{2(n-p)} \det(Q^T Q) \det(I_{n-p} + (Q^T Q)^{-1} Q^T Z D_\gamma Z^T Q) \\ &= \sigma^{2(n-p)} \det(Q^T Q) \det(I_q + D_\gamma Z^T Q (Q^T Q)^{-1} Q^T Z) \\ &= \sigma^{2(n-p)} \det(Q^T Q) \det(I_q + D_\gamma Z^T R Z), \end{aligned} \tag{23}$$

where

$$R = Q(Q^T Q)^{-1} Q^T = I_n - X(X^T X)^{-1} X^T = I_n - P_X.$$

From (17), $Q(Q^T V_\theta Q)^{-1} Q^T = R_\theta$. Then the REML log-likelihood function can also be expressed as

$$l_R(\theta) = -\frac{1}{2} \log\det(Q^T Q) - \frac{n-p}{2} \log(2\pi\sigma^2) - \frac{1}{2} \log\det(I_q + D_\gamma Z^T R Z) - \frac{1}{2} y^T R_\theta y. \tag{24}$$

This also shows that the REML estimator, obtained by maximizing $l_R(\theta)$, doesn't depend on the choice of $Q$. We can choose $Q$ such that $Q^T Q = I_{n-p}$. From the REML equations (18),

$$y^T R_{\hat{\theta}} V_{\hat{\theta}} R_{\hat{\theta}} y = tr(R_{\hat{\theta}} V_{\hat{\theta}}) = tr(I_n - V_{\hat{\theta}}^{-1} X (X^T V_{\hat{\theta}}^{-1} X)^{-1} X^T) = n - p.$$

From (8), $y^T R_{\hat{\theta}} y = y^T R_{\hat{\theta}} V_{\hat{\theta}} R_{\hat{\theta}} y = n - p$, and then the maximum REML log-likelihood function can be computed by

$$l_R(\hat{\theta}) = -\frac{n-p}{2} - \frac{n-p}{2} \log(2\pi\hat{\sigma}^2) - \frac{1}{2} \log\det(I_q + D_{\hat{\gamma}} Z^T R Z). \tag{25}$$

From $y^T R_\theta V_\theta R_\theta y = \sigma^{-2} y^T R_\gamma V_\gamma R_\gamma y$ and $V_\theta^{-1}(y - X\hat{\beta}) = R_\theta y$, we also have

$$\hat{\sigma}^2 = \frac{1}{n-p} y^T R_{\hat{\gamma}} V_{\hat{\gamma}} R_{\hat{\gamma}} y = \frac{1}{n-p} (y - X\hat{\beta})^T V_{\hat{\gamma}}^{-1} (y - X\hat{\beta}). \tag{26}$$

#### 1.2 Hypothesis testing

The hypotheses for testing fixed effects and variance components can be respectively defined as

$$H_{0,i} : \beta_i = 0 \ \text{ versus } \ H_{1,i} : \beta_i \neq 0,$$

$$H_{0,k} : \sigma_k^2 = 0 \ \text{ versus } \ H_{1,k} : \sigma_k^2 > 0.$$

Under regularity conditions, the MLE is consistent and asymptotically normal with asymptotic covariance matrix equal to the inverse of Fisher information matrix (Jiang 2007), that is, asymptotically

$$\hat{\beta} - \beta \sim N(\mathbf{0}, var(\hat{\beta})), \tag{27}$$

$$\hat{\theta} - \theta \sim N(\mathbf{0}, I(\theta)^{-1}). \tag{28}$$

Testing zero variance components is one of the most challenging problems because the null hypothesis, $\sigma_k^2 = 0$, is on the boundary of the parameter space (Drikvandi et al. 2012). The asymptotic normality is inappropriate for the null hypothesis with $\sigma_k^2 = 0$, and the usual asymptotic $\chi^2$ distribution of the likelihood ratio test (LRT) statistic under this null hypothesis is incorrect because the null is on the zero boundary of the parameter space. In this case, the asymptotic distribution of the LRT statistic is a mixture of $\chi^2$ distributions (Self and Liang (1987), Stram and Lee (1994)), which is determined by the degrees of freedom and the associated weights of the distributions in the mixture. Computing the degrees of freedom and the weights is computationally intensive.

To avoid the problem of zero-boundary in hypothesis testing, we reparameterize the variance components by $\theta_k = \sigma^2\gamma_k$, $k = 1, \ldots, K$, and allow the parameters, $\gamma_k$ or $\theta_k$, to take negative values. The lower boundary of the parameters can be a negative value such that the variance-covariance matrix

$$V_\theta = \sigma^2(I_n + \gamma_1 Z_1 Z_1^T + \ldots + \gamma_K Z_K Z_K^T),$$

is well defined (positive-definite). Note that $V_\theta$ is positive-definite when $\gamma_k > -1/\lambda_{max}$, where $\lambda_{max} > 0$, is the largest singular value of $ZZ^T$ or $Z^T Z$. That is, $V_\theta$ can be well-defined with negative parameters $\theta_k$. Then the hypotheses for the variance components are extended as

$$H_{0,k} : \theta_k \leq 0 \ \text{ versus } \ H_{1,k} : \theta_k > 0,$$

in which the zero components, $\theta_k = 0$, are no longer on the boundary of the parameter space, and the the MLE asymptotic properties hold under regularity conditions, which enables the use of z-statistics or t-statistics for testing fixed effects and variance components. If $\theta_k > 0$, $\sigma_k^2 = \theta_k$ is definable and the mixed-effects model is well-specified. Otherwise, $\theta_k \leq 0$, implies that the term of random effects is not needed in the model design.

The t-test statistics for fixed effects are given by

$$T_i = \frac{\hat{\beta}_i}{\sqrt{var(\hat{\beta}_i)}} = \frac{\hat{\beta}_i}{\sqrt{var(\hat{\beta})_{ii}}} \ \sim \ t(n-p). \tag{29}$$

The t-test statistic for a contrast, a linear combination of the estimated fixed effects, $c^T\hat{\beta}$, is

$$T_c = \frac{c^T\hat{\beta}}{\sqrt{c^T var(\hat{\beta}) c}} \sim t(n-p). \tag{30}$$

The z-test statistics for the parameters of variance components are given by

$$Z_{\theta_k} = \frac{\hat{\theta}_k}{\sqrt{[I(\hat{\theta})^{-1}]_{kk}}} \sim N(0,1). \tag{31}$$

If $Z_{\theta_k} > 0$, then $\sigma_k^2 = \theta_k$ is definable. Otherwise, $Z_{\theta_k} \leq 0$, means that the random effects are not required or can be ignored.

We can also use likelihood ratio test (LRT) statistic to test the variance components. Let $\theta_{sub}$ be the sub-vector of $\theta$ that are not equal to zero. Let $L(\hat{\theta}_{sub})$ and $L(\hat{\theta})$ be the maximum likelihood for the sub-model and the full model, respectively. Note that $L(\hat{\theta}_{sub})$ and $L(\hat{\theta})$ depend on $\hat{\beta}$ for ML method which is suppressed for simplicity of notation. The LRT statistic,

$$\lambda_{LR} = 2[\log(L(\hat{\theta})) - \log(L(\hat{\theta}_{sub}))] = 2[l(\hat{\theta}) - l(\hat{\theta}_{sub})] \sim \chi_r^2, \tag{32}$$

a asymptotic $\chi^2$ distribution, where $r$ is the number of zero variance components. Note that, for REML method, the fixed-effects design matrix should be the same for both sub and full models.

#### 1.3 Prediction of random effects

With the assumption of normality: $b \sim N(\mathbf{0}, D_\theta)$ and $\epsilon \sim N(\mathbf{0}, \sigma^2 I_n)$, from (1), we have

$$\begin{bmatrix} b \\ y \end{bmatrix} \sim N\Big( \begin{bmatrix} \mathbf{0} \\ X\beta \end{bmatrix}, \begin{bmatrix} D_\theta & D_\theta Z^T \\ ZD_\theta & V_\theta \end{bmatrix} \Big),$$

Note $V_\theta = \sigma^2 I_n + ZD_\theta Z^T$. The prediction of random effects is given by a conditional mean:

$$\hat{b} = E(b|y) = D_\theta Z^T V_\theta^{-1}(y - X\beta). \tag{33}$$

Since

$$E_{b,y}(\| \hat{b} - b \|^2) = E_y(\| \hat{b} - E(b|y) \|^2) + E_{b,y}(\| b - E(b|y) \|^2),$$

$\hat{b} = E(b|y)$, is the best predictor in the sense of minimum mean squared error (MSE) of prediction . It is seen from $E(\hat{b}) = E_y[E(b|y)] = E(b)$, that the best predictor is unbiased. Without any assumption of normality, the prediction (33) can also be derived from the best linear predictor of the form

$$b_{BLP} = a + A[y - E(y)],$$

by minimizing $E(\| b_{BLP} - b \|^2)$ which yields

$$a = E(b) = \mathbf{0}, \quad A = Cov(b, y)[Cov(y)]^{-1} = D_\theta Z^T V_\theta^{-1}.$$

Therefore the $\hat{b}$ is the best linear unbiased prediction (BLUP) (Searle, Casella, and McCulloch 2006). It is also known that the MLE of $\beta$,

$$\hat{\beta} = (X^T V_\theta^{-1} X)^{-1} X^T V_\theta^{-1} y,$$

is the best linear unbiased estimator (BLUE), which does not require the normality assumption. This can be readily verified. Let $\beta_{LUE} = \hat{\beta} + Cy$ be any linear unbiased estimator of $\beta$. Then we have $CX = \mathbf{0}$ and

$$E(\| \beta_{LUE} - \beta \|^2) = E(\| \hat{\beta} - \beta \|^2) + tr(CV_\theta C^T) \geq E(\| \hat{\beta} - \beta \|^2).$$

Thus $\hat{\beta}$ minimizing the MSE is the BLUE.

The BLUP of $E(y|b) = X\beta + Zb$ is give by

$$\begin{aligned} \hat{y} &= X\beta + Z\hat{b} \\ &= X\beta + ZD_\theta Z^T V_\theta^{-1}(y - X\beta) \\ &= X\beta + (I_n - \sigma^2 V_\theta^{-1})(y - X\beta) \\ &= y - \sigma^2 V_\theta^{-1}(y - X\beta). \end{aligned} \tag{34}$$

Substituting $\hat{\beta}$ and $\hat{\theta}$ for the unknown parameters in (33) and (34), we have the empirical BLUPs (Harville 1991),

$$\hat{b} = E(b|y) = D_{\hat{\theta}} Z^T V_{\hat{\theta}}^{-1}(y - X\hat{\beta}) = D_{\hat{\theta}} Z^T R_{\hat{\theta}} y, \tag{35}$$

$$\hat{y} = E(y|b) = y - \hat{\sigma}^2 R_{\hat{\theta}} y, \tag{36}$$

where $R_\theta$ is defined by (6), that is,

$$R_\theta = V_\theta^{-1} - V_\theta^{-1} X (X^T V_\theta^{-1} X)^{-1} X^T V_\theta^{-1}.$$

##### 1.4 Numerical algorithms

With variance components estimated, the fixed effects estimated by ML and REML methods are given by (7) or (19) as follows

$$\hat{\beta} = (X^T V_\theta^{-1} X)^{-1} X^T V_\theta^{-1} y.$$

Estimating variance components by either ML or REML is a numerical optimization problem. Various iterative methods based on the log likelihood, called gradient methods, have been proposed to compute the ML and REML estimates (Searle, Casella, and McCulloch 2006). The gradient methods are represented by the iteration equation

$$\theta^{(i+1)} = \theta^{(i)} + \Gamma(\theta^{(i)}) \frac{\partial l(\theta^{(i)})}{\partial \theta}, \tag{37}$$

where $\theta^{(i)}$ represents the value of the estimate of the parameter vector $\theta$ at the $i$th iteration, $\partial l(\theta)/\partial\theta$ is the gradient of the log likelihood function, and $\Gamma(\theta)$ is a modifier matrix of the gradient direction. Let $H(\theta)$ and $I(\theta)$ be the Hessian matrix and Fisher information matrix of the log likelihood function with respect to $\theta$. Let $I_A(\theta) = [I(\theta) - H(\theta)]/2$ be the average information matrix. The modifier matrix can be specified by

1) Newton–Raphson: $\Gamma(\theta) = -H(\theta)^{-1}$;
2) Fisher scoring: $\Gamma(\theta) = I(\theta)^{-1}$;
3) Average information: $\Gamma(\theta) = I_A(\theta)^{-1}$.

For MLE, from (4), (5), (9), (10), and $V_\theta^{-1}(y - X\hat{\beta}) = R_\theta y$, we have

$$\begin{array}{l} \frac{\partial l}{\partial \theta_i} = -\frac{1}{2}[tr(V_\theta^{-1} V_i) - y^T R_\theta V_i R_\theta y], \\ H_{ij} = \frac{\partial^2 l}{\partial\theta_i \partial\theta_j}|_{\theta,\hat{\beta}} = \frac{1}{2} tr(V_\theta^{-1} V_i V_\theta^{-1} V_j) - y^T R_\theta V_i V_\theta^{-1} V_j R_\theta y, \\ I_{ij} = -E(\frac{\partial^2 l}{\partial\theta_i\partial\theta_j}) = \frac{1}{2} tr(V_\theta^{-1} V_i V_\theta^{-1} V_j), \\ I_{A,ij} = \frac{1}{2}(I_{ij} - H_{ij}) = \frac{1}{2} y^T R_\theta V_i V_\theta^{-1} V_j R_\theta y. \end{array} \tag{38}$$

For REML, $l = l_R(\theta)$. From (20), (21) and (22), we have

$$\begin{array}{l} \frac{\partial l_R}{\partial \theta_i} = -\frac{1}{2}[tr(R_\theta V_i) - y^T R_\theta V_i R_\theta y], \\ H_{ij} = \frac{\partial^2 l_R}{\partial\theta_i \partial\theta_j} = \frac{1}{2} tr(R_\theta V_i R_\theta V_j) - y^T R_\theta V_i R_\theta V_j R_\theta y, \\ I_{ij} = -E(H_{ij}) = \frac{1}{2} tr(R_\theta V_i R_\theta V_j), \\ I_{A,ij} = \frac{1}{2}(I_{ij} - H_{ij}) = \frac{1}{2} y^T R_\theta V_i R_\theta V_j R_\theta y. \end{array} \tag{39}$$

Here $H_{ij}$, $I_{ij}$ and $I_{A,ij}$ represent the $(i,j)$ entry of $H(\theta)$, $I(\theta)$ and $I_A(\theta)$, respectively. Recall $V_i = Z_i Z_i^T$, defined in (4), and

$$V_\theta = \sigma^2 I_n + Z D_\theta Z^T,$$

$$R_\theta = V_\theta^{-1} - V_\theta^{-1} X (X^T V_\theta^{-1} X)^{-1} X^T V_\theta^{-1},$$

defined in (2) and by (6), respectively.

#### 2 Summary statistics based algorithm

This section describes our work to improve LMM solver scalability and memory usage efficiency in the context of single cell transcriptomics data. The gradient methods given by (37), (38) and (39) have a computational complexity of $O(n^3)$. The computational burden arises from the $n \times n$ matrix inverse, $V^{-1}(\theta)$, and the matrix-matrix products, $V_\theta^{-1} V_i V_\theta^{-1} V_j$ and $R_\theta V_i R_\theta V_j$. Now we derive a summary statistics based algorithm to implement the gradient methods for the LMM estimation. The $n \times n$ matrix inverse and matrix-matrix products are computed through the low dimension $p \times p$ and $q \times q$ matrices. The algorithm achieves a computational complexity of $O(n(p^2 + q^2))$, which is fast and linearly scalable with the sample size $n$. In addition, using the summary statistics requires less computer memory usage to estimate the LMM parameters, which enables the LMM application to the large-scale data analysis. By precomputing and directly using the summary statistics as inputs, the algorithm has a computational complexity of $O(p^3+q^3)$, which makes computations independent of the sample size $n$ and achieves both speed and memory efficiency.

Recall that $\gamma_k = \sigma_k^2/\sigma^2$, $k=1,\ldots,K$, and

$$V_\gamma = I_n + \gamma_1 Z_1 Z_1^T + \ldots + \gamma_K Z_K Z_K^T = I_n + ZDZ^T,$$

$$R_\gamma = V_\gamma^{-1} - V_\gamma^{-1}X(X^T V_\gamma^{-1} X)^{-1} X^T V_\gamma^{-1},$$

where $D = D_\gamma$. Then $V_\theta = \sigma^2 V_\gamma$, $R_\theta = \sigma^{-2} R_\gamma$, and

$$\begin{array}{rcl} tr(V_\theta^{-1}V_i) & = & \sigma^{-2} tr(Z_i^T V_\gamma^{-1} Z_i), \\ tr(V_\theta^{-1}V_i V_\theta^{-1} V_j) & = & \sigma^{-4} tr[(Z_i^T V_\gamma^{-1} Z_j)^T (Z_i^T V_\gamma^{-1} Z_j)], \\ y^T R_\theta V_i V_\theta^{-1} V_j R_\theta y & = & \sigma^{-6} tr[(Z_i^T R_\gamma y)^T (Z_i^T V_\gamma^{-1} Z_j)(Z_j^T R_\gamma y)], \\ tr(R_\theta V_i) & = & \sigma^{-2} tr(Z_i^T R_\gamma Z_i), \\ tr(R_\theta V_i R_\theta V_j) & = & \sigma^{-4} tr[(Z_i^T R_\gamma Z_j)^T (Z_i^T R_\gamma Z_j)], \\ y^T R_\theta V_i R_\theta y & = & \sigma^{-4} tr[(Z_i^T R_\gamma y)^T (Z_i^T R_\gamma y)], \\ y^T R_\theta V_i R_\theta V_j R_\theta y & = & \sigma^{-6} tr[(Z_i^T R_\gamma y)^T (Z_i^T R_\gamma Z_j)(Z_j^T R_\gamma y)]. \end{array} \tag{40}$$

Using the formulas of matrix identities given by Harville (1977) and Rao (1979), we have

$$V_\gamma^{-1} = I_n - Z(I_q + DZ^T Z)^{-1} DZ^T,$$

$$R_\gamma = R - RZ(I_q + DZ^T RZ)^{-1} DZ^T R,$$

where

$$R = I_n - X(X^T X)^{-1} X^T.$$

Let

$$\begin{array}{rcl} M_0 & = & (I_q + DZ^T Z)^{-1}, \\ M & = & (I_q + DZ^T RZ)^{-1}. \end{array} \tag{41}$$

Then $V_\gamma^{-1} = I_n - ZM_0DZ^T$ and $R_\gamma = R - RZMDZ^T R$. By the identity equation

$$DZ^T Z M_0 = M_0 D Z^T Z = I_q - M_0,$$

$$DZ^T RZM = MDZ^T RZ = I_q - M,$$

and $R^2 = R$, we have $V_\gamma^{-2} = V_\gamma^{-1} - ZM_0^2 DZ^T$, $R_\gamma^2 = R_\gamma - RZM^2DZ^T R$, and

$$\begin{array}{rcl} Z_i^T V_\gamma^{-1} Z_j & = & Z_i^T Z_j - (Z_i^T Z) M_0 D (Z^T Z_j), \\ Z_i^T V_\gamma^{-2} Z_i & = & Z_i^T V_\gamma^{-1} Z_i - (Z_i^T Z) M_0^2 D (Z^T Z_i), \\ tr(V_\gamma^{-1}) & = & n - q + tr(M_0), \\ tr(V_\gamma^{-2}) & = & n - q + tr(M_0^2), \end{array} \tag{42}$$

$$\begin{array}{rcl} Z_i^T R_\gamma Z_j & = & Z_i^T RZ_j - (Z_i^T RZ) MD (Z^T RZ_j), \\ Z_i^T R_\gamma^2 Z_i & = & Z_i^T R_\gamma Z_i - (Z_i^T RZ) M^2 D (Z^T RZ_i), \\ tr(R_\gamma) & = & tr(R) - q + tr(M), \\ tr(R_\gamma^2) & = & tr(R) - q + tr(M^2), \end{array} \tag{43}$$

$$\begin{array}{rcl} Z_i^T R_\gamma y & = & Z_i^T Ry - (Z_i^T RZ) MD (Z^T Ry), \\ y^T R_\gamma^2 y & = & y^T R_\gamma y - (y^T RZ) M^2 D (Z^T Ry), \\ y^T R_\gamma y & = & y^T Ry - (y^T RZ) MD (Z^T Ry), \end{array} \tag{44}$$

where

$$\begin{array}{c} Z_i^T RZ_j = Z_i^T Z_j - (Z_i^T X)(X^T X)^{-1}(X^T Z_j), \\ Z^T Ry = Z^T y - (Z^T X)(X^T X)^{-1}(X^T y), \\ y^T Ry = y^T y - (y^T X)(X^T X)^{-1}(X^T y). \end{array} \tag{45}$$

From the formulas (40), (42), (43) and (44), $V_i = Z_i Z_i^T$, $V_\theta^{-1} = \sigma^{-2} V_\gamma^{-1}$, and $R_\theta = \sigma^{-2} R_\gamma$, we have the following formulas, which are used to compute the gradients, $\frac{\partial l}{\partial \theta_i}$ and $\frac{\partial l_R}{\partial \theta_i}$, given by (38) and (39),

$$\begin{aligned} tr(V_\theta^{-1} V_i) &= \sigma^{-2}[tr(Z_i^T Z_i) - tr((Z_i^T Z) M_0 D (Z^T Z_i))], \\ tr(R_\theta V_i) &= \sigma^{-2}[tr(Z_i^T R Z_i) - tr((Z_i^T R Z) M D (Z^T R Z_i))], \\ y^T R_\theta V_i R_\theta y &= \sigma^{-4} tr\{[Z_i^T R y - (Z_i^T R Z) M D (Z^T R y)]^T [Z_i^T R y - (Z_i^T R Z) M D (Z^T R y)]\} \end{aligned} \tag{46}$$

Thus the gradient methods given by (37), (38) and (39) can be implemented by using the summary statistics: $X^T X$, $X^T Z$, $X^T y$, $Z^T y$ and $Z^T Z$, based on the formulas (40)~(46).

#### 2.1 Computational complexity

The complexity for computing the summary statistics: $X^T X$, $X^T y$, $Z^T X$, $Z^T y$ and $Z^T Z$, is $O(n(p^2 + q^2))$. The complexity for computing the $q \times q$ matrix inverse, $M_0$ and $M$, by (41) is $O(q^3)$ or even $O(q^2 \log(q))$. Using the summary statistics as the inputs, the algorithms given by (43), (44) and (45) have a complexity of $O(p^3 + q^3)$. Then the summary statistics based algorithm given by equations (40)~(45) has the complexity

$$O(n(p^2 + q^2) + p^3 + q^3) = O(n(p^2 + q^2)),$$

since $p < n$, $q_k < n$ and $q < Kn$. The summary statistics can be computed in advance and stored in a computer with less storage. Once the summary statistics are computed and directly used as the inputs, the algorithm complexity, $O(p^3 + q^3)$, doesn't depend on the sample size $n$.

#### 2.2 LMM with dimension reduction

The LMM estimation algorithm can be further sped up by reducing the number of random effects. For a large number of random effects, we may combine the correlated random effects by cluster analysis or principal component analysis (PCA). By PCA, we have

$$Z_k = U_k V_k^T,$$

where $U_k$ is an $n \times r_k$ matrix of principal components (PCs), $V_k$ is an $q_k \times r_k$ matrix, $r_k \leq \min(n, q_k)$ is the rank of $Z_k$, $U_k^T U_k$ is diagonal, and $V_k^T V_k = I_{r_k}$, $k = 1, \ldots, K$. Thus the LMM (1) can be rewritten as

$$y = X\beta + U_1 v_1 + \ldots + U_K v_K + \epsilon = X\beta + Uv + \epsilon, \tag{47}$$

where $v_k = V_k^T b_k \sim N(\mathbf{0}, \sigma_k^2 I_{r_k})$, is a $r_k$-vector of random effects. The LMM (47) is equivalent to the LMM (1). The random effect design matrices $U_k$ in the LMM (47) have a lower dimension $r_k$. We may even approximate the design matrix $Z_k$ using a smaller number of PCs such that $r_k \ll \min(n, q_k)$.

### 3 Simulation studies on the LMM hypothesis testing

We examined the performance of the FLASH-MM t-test, z-test and likelihood ratio test (LRT), given by (29), (31) and (32), in controlling Type I error rate by comparing

- the Satterthwaite approximation's t-test for fixed effects, implemented in lmerTest package (Kuznetsova, Brockhoff, and Christensen 2017);
- lme4 package anova (Bates et al. 2015) and varTestnlme package LRT (Baey and Kuhn 2023) for testing variance components.

The lmerTest package provides tools for testing fixed effects. The varTestnlme package allows testing whether a subset of the variance components is equal to zero, using the asymptotic properties of the LRT statistic. The varTestnlme requires the model to be fit by maximum likelihood (ML) rather than restricted maximum likelihood (REML).

We considered the LMM with random intercepts and random slopes:

$$y_{ij} = (\beta_1 + \beta_2 x_{ij}) + (b_{1i} + b_{2i} x_{ij}) + \epsilon_{ij}, \tag{48}$$

where $x_{ij}$, $i = 1, \dots, N$ and $j = 1, \dots, n_i$, is the $j$th observation of a covariate for the $i$th individual, $\beta_1$ and $\beta_2$ are the fixed effects for the intercept and slope, respectively, and $b_{1i}$ and $b_{2i}$ are the random intercept and slope, respectively. $b_{1i}$, $b_{2i}$ and $\epsilon_{ij}$ are independent and normally distributed with zero mean and variances $\sigma_1^2$, $\sigma_2^2$, and $\sigma^2$, respectively. Let $x_{ij} = 1 + j/n_i$. We generated 1000 simulation samples from the model (48) with $N = 25$ and $n_i$ randomly generated by the Poisson function $rpois(N, lambda)$ with $lambda = 5,\ 10,\ 25$ and 50, respectively. The simulated data are unbalanced. The $lambda$ is an average number of the observations per individual.

**Case 1**: Testing the fixed effect of the slope:

$$H_0 : \beta_2 = 0 \ \text{ versus } \ H_1 : \beta_2 \neq 0.$$

In this case, $\beta_1 = 1$, $\beta_2 = 0$, $\sigma_1^2 = \sigma_2^2 = 0.25$, and $\sigma^2 = 1$. The QQ-plots of the FLASH-MM and lmerTest p-values for testing the slope $\beta_2 = 0$ are shown in Figure S1. It is seen that FLASH-MM t-test has a good Type-I error control as lmerTest Satterthwaite approximation's t-test because the points in the QQ-plots fall within the 95% confidence band.

**Case 2**: Testing the variance of the random slope:

$$H_0 : \sigma_2^2 = 0 \ \text{ versus } \ H_1 : \sigma_2^2 \neq 0.$$

In this case, $\beta_1 = \beta_2 = 1$, $\sigma_1^2 = 0.25$, $\sigma_2^2 = 0$, and $\sigma^2 = 1$. The QQ-plots of the FLASH-MM z-test and LRT, varTestnlme LRT and lme4 anova p-values for testing the variance of random slope $\sigma_2^2 = 0$ are shown in Figure S2. It is seen that FLASH-MM z-test and LRT have a good Type-I error control. The lme4 anova deflates the p-values. The varTestnlme LRT deflates the p-values for small number of observations ($lambda = 5$ and 10). As the number of observations increases ($lambda = 25$ and 50), both FLASH-MM LRT and varTestnlme LRT have a similar performance.

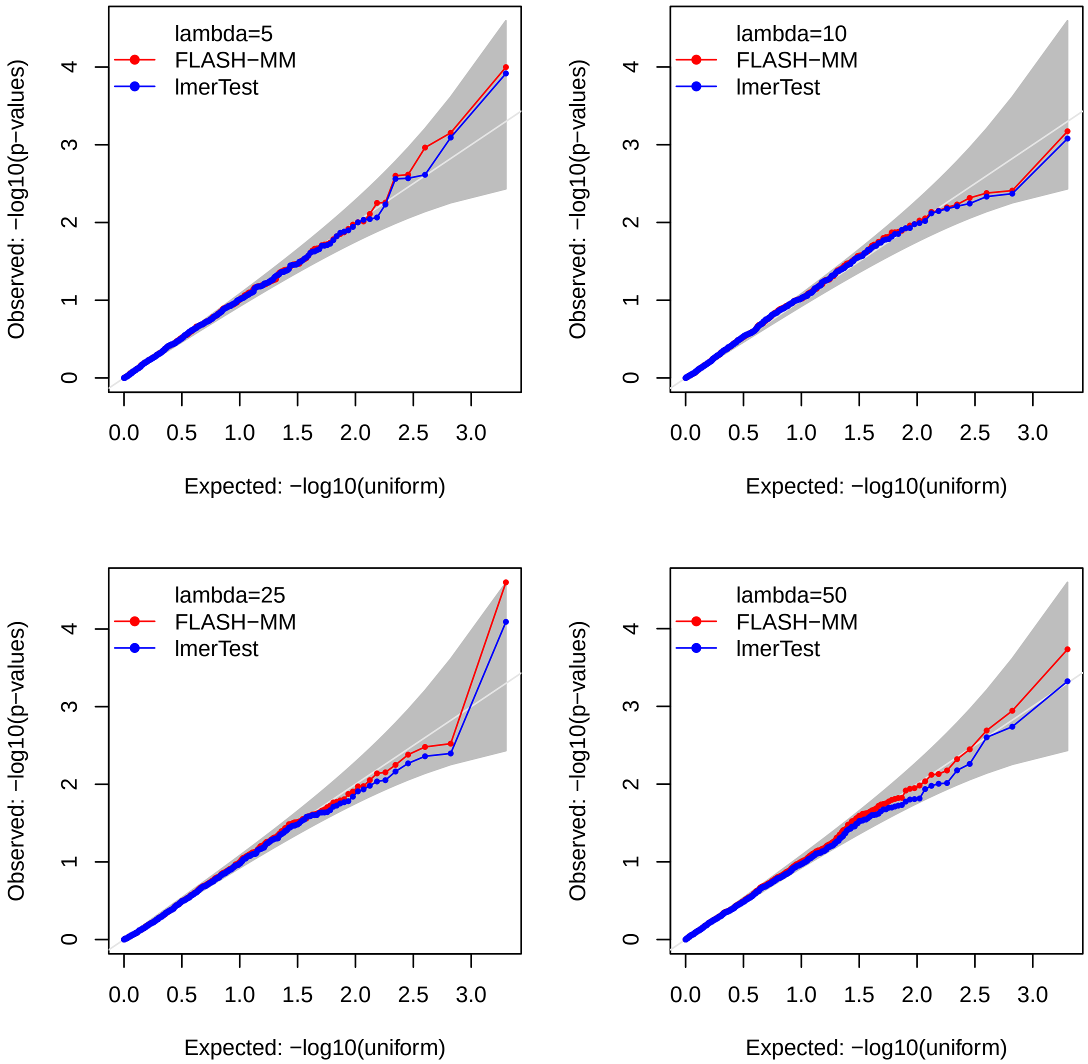

Figure S1: QQ-plots of the FLASH-MM and lmerTest p-values for testing the slope $\beta_2 = 0$ in the models with $lambda = 5,\ 10,\ 25$ and 50, respectively.

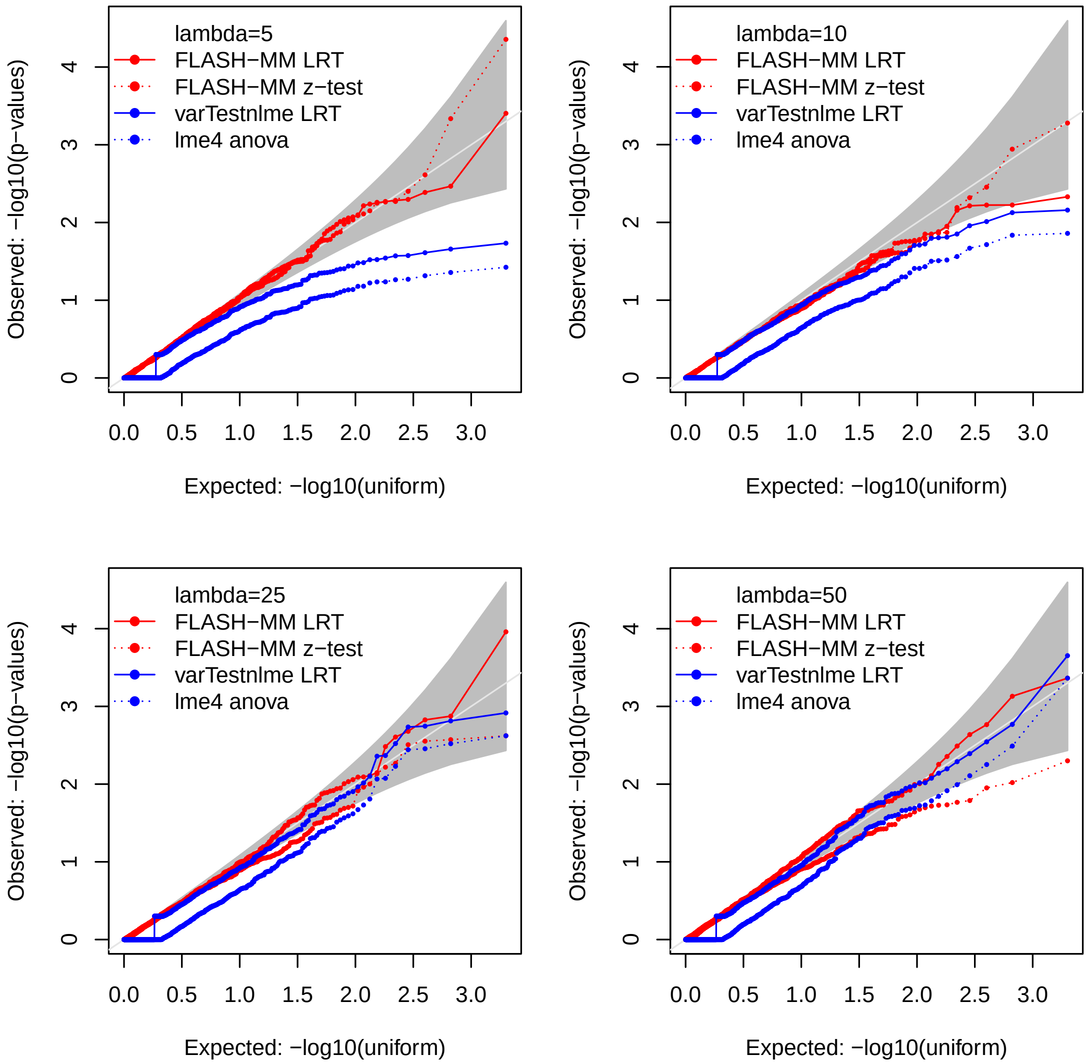

Figure S2: QQ-plots of the FLASH-MM z-test, varTestnlme LRT and lme4 anova p-values for testing the variance of the random slope $\sigma_2^2 = 0$ in the models with $lambda = 5,\ 10,\ 25$ and 50, respectively.

### 4 Simulation studies in scRNA-seq differential expression analysis

#### 4.1 Simulation methods

To simulate the multi-subject multi-cell-type single-cell RNA-seq (scRNA-seq) dataset, we developed a scRNA-seq simulator, named simuRNAseq, by using a reference dataset based on a negative binomial (NB) distribution. The reference dataset contains genes-by-cells counts matrix and meta data. The simulated genes are randomly selected from the reference data. If the number of simulated genes is equal to the number of genes in the reference data, the original gene names in the reference data are retained. The simulated cells are randomly selected from the meta data that specifies subjects, cell-types and treatment conditions for the cells. If the reference data is not available, it will be generated randomly. The simuRNAseq workflow is illustrated in the Supplemental Figure S3, which consists of following main steps:

(1) Estimate the dispersion and mean of the NB distribution for each gene using the reference count data.
(2) Select the differetially expressed (DE) genes and non-DE genes randomly from the genes in the reference

count data and the cells from the meta data, with given numbers.

(3) Generate counts by sampling from the NB distributions for the non-DE genes without log-fold-change (logFC) between treatments and for the DE genes with assigned logFC between treatments.

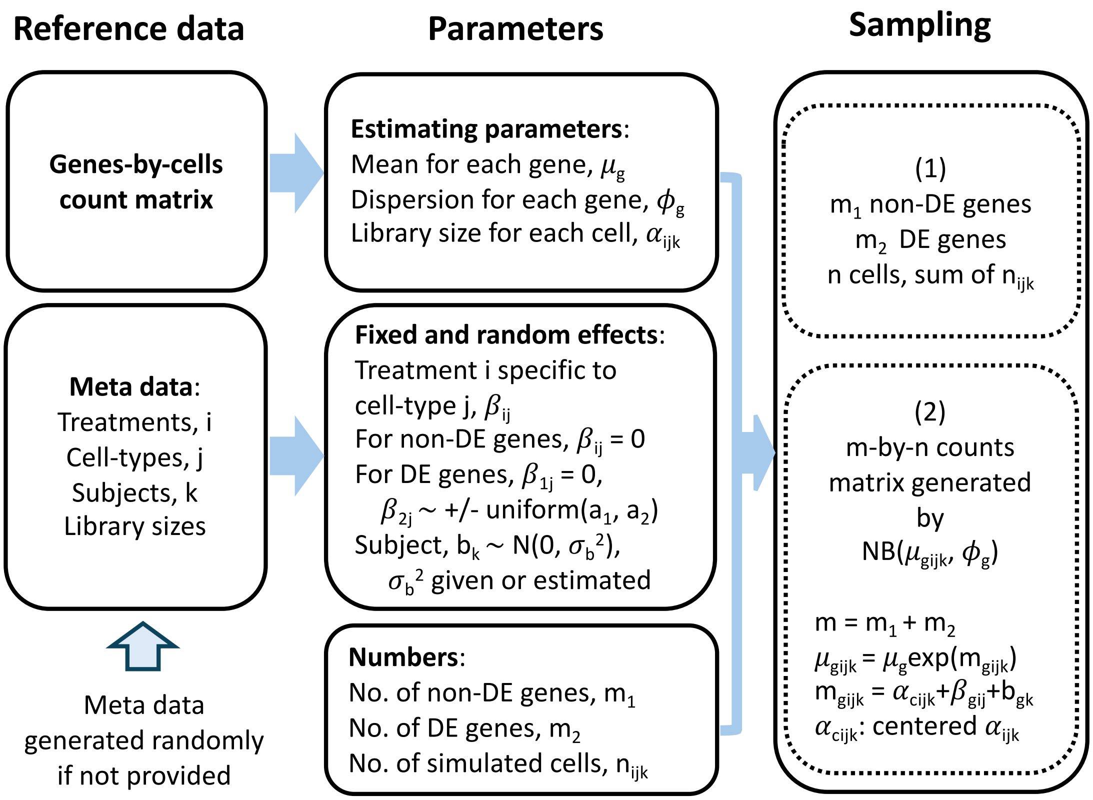

Figure S3: Workflow of the scRNA-seq simulator, simuRNAseq.

Let $y_{g,ijk}$ be the count for gene $g$ and the cell from subject $k = 1, \ldots, n_s$ and cell-type $j = 1, \ldots, n_c$ with treatment $i = 1, 2$. The count is generated by the NB distribution with dispersion $\phi_g$ and mean $\mu_{g,ijk}$,

$$y_{g,ijk} \sim NB(\mu_{g,ijk}, \phi_g), \tag{49}$$

where

$$\log(\mu_{g,ijk}) = \log(\mu_g) + \alpha_{ijk} + \beta_{g,ij} + b_{g,k},$$

$\phi_g$ and $\mu_g$ are the dispersion and mean estimated from the reference data for gene $g$, $\alpha_{ijk}$ is a centered log-library-size for the cell $c_{ijk}$, $\beta_{g,ij}$ represents the effect of treatment $i$ specific to cell-type $j$, $b_{g,k} \sim N(0, \sigma_b^2)$ is a random effect of subject variation. For the non-DE genes, $\beta_{g,1j} = \beta_{g,2j} = 0$. For the DE genes, $\beta_{g,1j} = 0$ and $\beta_{g,2j} \sim \pm Uniform(a_1, a_2)$, a uniform distribution with $a_1 a_2 > 0$, where $a_1$ and $a_2$ are the lower and upper bounds of the DE gene effect sizes.

The mean of NB distribution is taken as the sample mean for each gene. The dispersion of NB distribution is computed by the method-of-moments estimate (MME) (Clark and Perry 1989), which is computationally more simple and performs reasonably well compared to the maximum likelihood estimate (MLE). The simuRNAseq simulator shares similarities with muscat (Crowell et al. 2020) and GLMsim (Wang et al. 2024). However, both muscat and GLMsim have limitations in scRNA-seq simulations. Muscat estimates the dispersion of the negative binomial (NB) distribution using the edgeR package based on a subset of the reference data. Muscat cannot be applied to large-scale scRNA-seq data and only reflects partial information from the reference dataset. GLMsim estimates the coefficients and dispersion parameters of the NB models for each gene using glm.nb from the MASS package (Venables and Ripley 2002), which is time-consuming and only generates data of a fixed size matching the reference data. In contrast, The simuRNAseq simulator is fast and flexible, and utilizes the whole information from the reference dataset.

We used the PBMC scRNA-seq data as a reference to simulate the scRNA-seq data. The PBMC data from eight lupus patients (Kang et al. 2018), available through Bioconductor's ExperimentHub package, contains 35,635 genes and 29,065 cells (14,619 control cells and 14,446 stimulated cells) consisting of eight identified cell types. After quality control filtering (filtering the cells with few or many detected genes and the genes lowly expressed), the data contains 7,040 genes and 26,820 cells.

#### 4.2 Performance of simuRNAseq simulator

We examined the performance of simuRNAseq simulator by comparing dispersions estimated by MME and MLE, means and variances across cells and library sizes of real reference data and simulated data using scatterplots. The simulated scRNA-seq data is generated by simuRNAseq with the same genes and cells as the reference data.

It is shown that the dispersions estimated by MME and MLE methods are coincident, see Supplemental Figure S4 (a). The MME method took only 0.48 seconds while the MLE method needed about 51.5 minutes to estimate the dispersions. This verified that the MME method is fast and performs very well. The means and variances of counts across cells and library sizes of the real and simulated data are shown in Supplemental Figure S4 (b), (c) and (d), respectively. The scatterplots display the similarity between the real and simulated data.

#### 4.3 Differential expression analysis of simulated scRNA-seq data

We generated scRNA-seq datasets consisting of 6,000 genes with 6 different numbers of cells (sample sizes) from 20,000 to 120,000 using the PBMC data as a reference, as described above. The genes to be simulated were randomly selected from the reference data. We randomly generated the meta data comprising 25 subjects and 12 cell-types which were treated by one of two treatments. The treatments, cell-types and subjects are assigned randomly with equal probability. There are 480 DE genes specific to a cell-type. For the DE genes $b_{g,2j} \sim \pm Uniform(0.25, 1)$. The variance component $\sigma_b^2 = 0.16$.

We did differential gene expression analysis for the 6 simulated scRNA-seq datasets using LMM with model formula:

$$\sim \log(library.size) + cell.type + cell.type : treatment + (1|subject). \tag{50}$$

The interaction term $cell.type : treatment$ in the model formula (50) represents treatment effects in a specific cell-type. The last term represent random effects of subject variation. We fit the LMM to the log-transformed counts, $log_2(1 + y)$, by FLASH-MM method. Note that FLASH-MM doesn't directly use model formula as an argument and need design matrices of fixed and random effects that can be created by, respectively,

$$X = model.matrix(\sim log(library.size) + cell.type + cell.type : treatment),$$

$$Z = model.matrix(\sim 0 + subject).$$

Table S1: Computation time in minutes for running FLASH-MM, lmer and nebula in differential expression analysis of the simulated data across various sample sizes $n$.

| | n=20000 | n=40000 | n=60000 | n=80000 | n=100000 | n=120000 |
|---|---|---|---|---|---|---|
| FLASH-MM | 0.46 | 0.51 | 0.67 | 0.67 | 0.88 | 1.04 |
| lmer | 25.15 | 49.70 | 74.90 | 94.02 | 122.44 | 149.12 |
| nebula | 39.71 | 86.45 | 140.95 | 205.98 | 274.87 | 331.89 |

Searle, Shayle R., George Casella, and Charles E. McCulloch. 2006. *Variance Components.* New Jersey: John Wiley & Sons, Inc.

Self, Steven G., and Kung-Yee Liang. 1987. "Asymptotic Properties of Maximum Likelihood Estimators and Likelihood Ratio Tests Under Nonstandard Conditions." *Journal of the American Statistical Association* 82 (398): 605–10.

Stram, Daniel O., and Jae Won Lee. 1994. "Variance Components Testing in the Longitudinal Mixed Effects Model." *Biometrics* 50 (4): 1171–77.

Venables, W. N., and B. D. Ripley. 2002. *Modern Applied Statistics with s.* Fourth. New York: Springer. https://www.stats.ox.ac.uk/pub/MASS4/.

Wang, Jianan, Lizhong Chen, Rachel Thijssen, Belinda Phipson, and Terence P. Speed. 2024. "GLMsim: A GLM-Based Single Cell RNA-Seq Simulator Incorporating Batch and Biological Effects." *bioRxiv.* https://doi.org/10.1101/2024.03.20.586030.

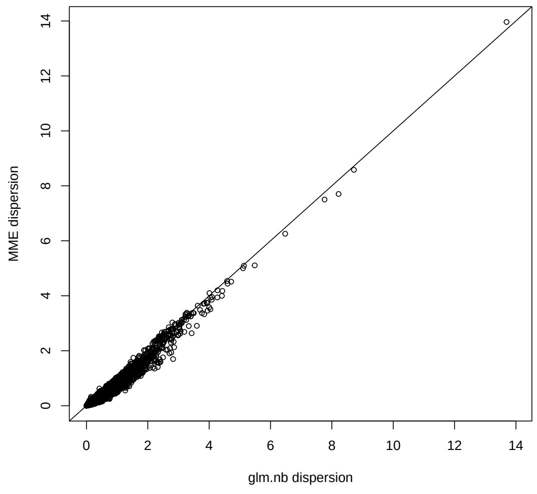

(a) Dispersions

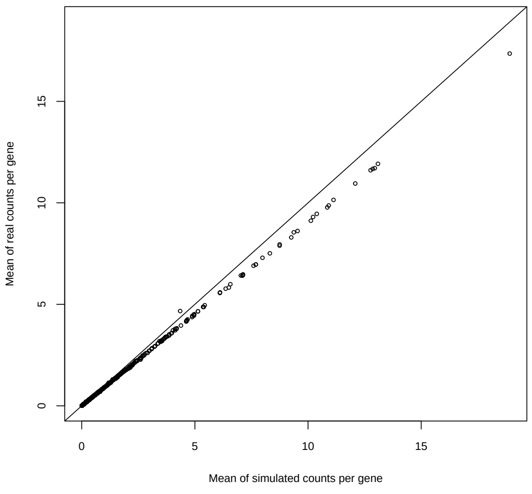

(b) Means

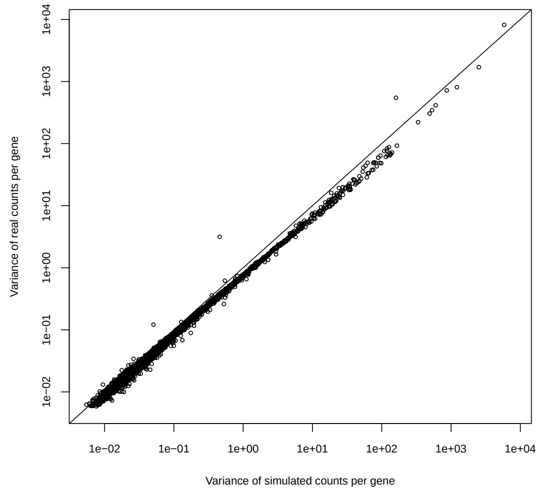

(c) Variances

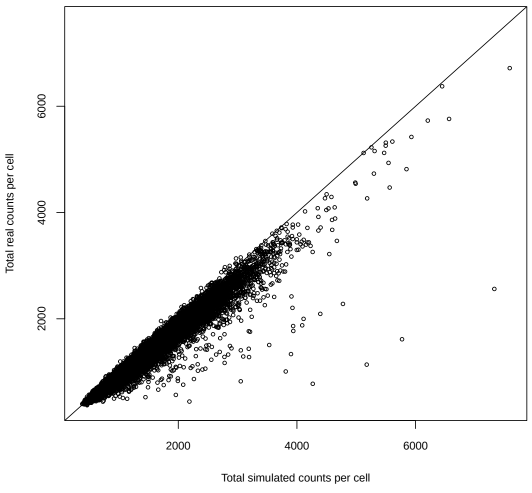

(d) Library sizes

Figure S4: **Performance of the scRNA-seq simulator, simuRNAseq**: (a) Scatterplot of dispersions estimated by MME and MLE (glm.nb). (b) Scatterplot of means of real and simulated counts across cells. (c) Scatterplot of variances of real and simulated counts across cells. (d) Scatterplot of library sizes of real and simulated counts.
