## Extended data file for "FLASH-MM: fast and scalable single-cell differential expression analysis using linear mixed-effects models"

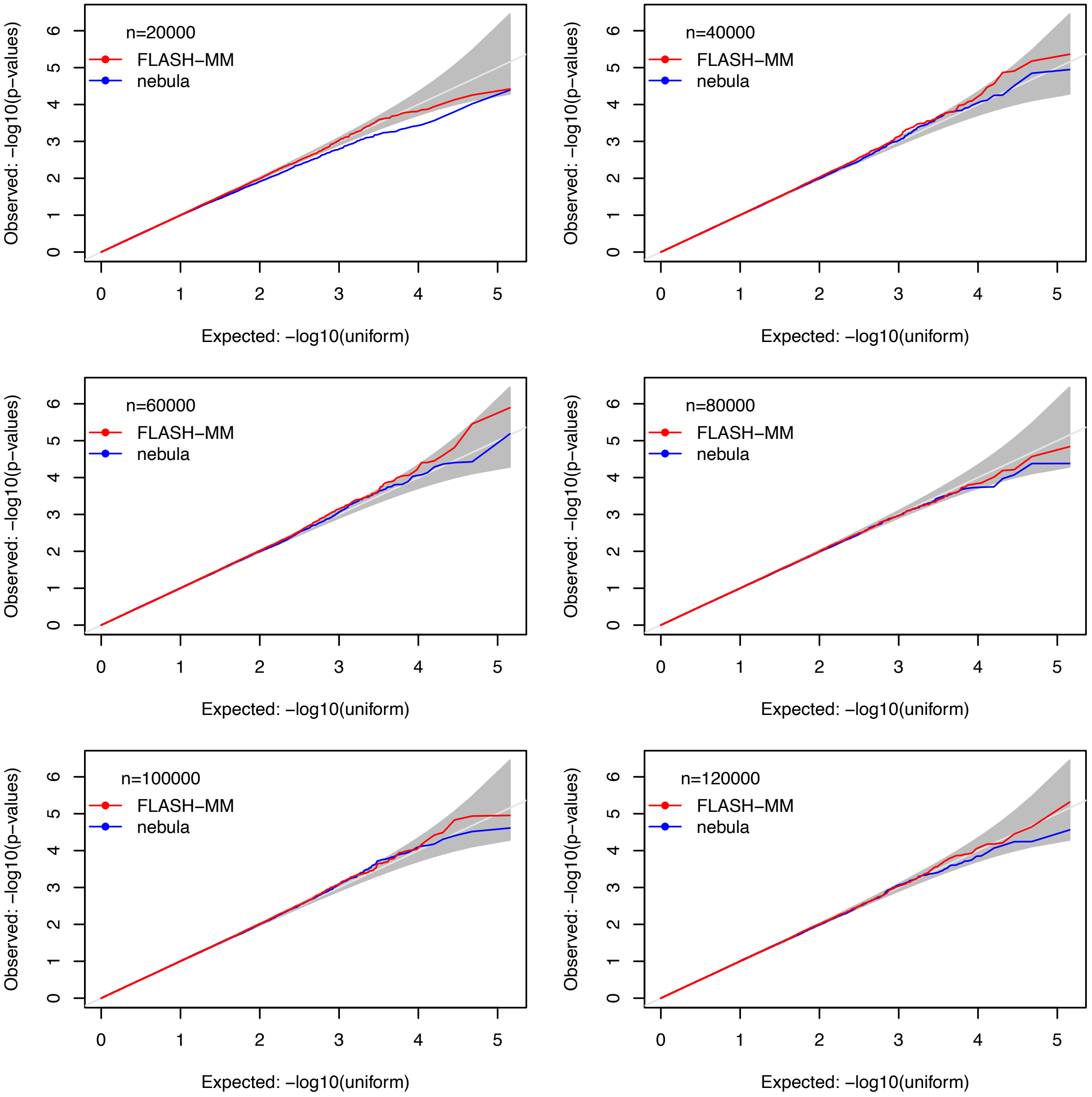


Extended Figure 1. QQ-plots of FLASH-MM and nebula p-values. QQ-plots of FLASH-MM and nebula p-values under the null hypothesis H0 in different sample sizes n.

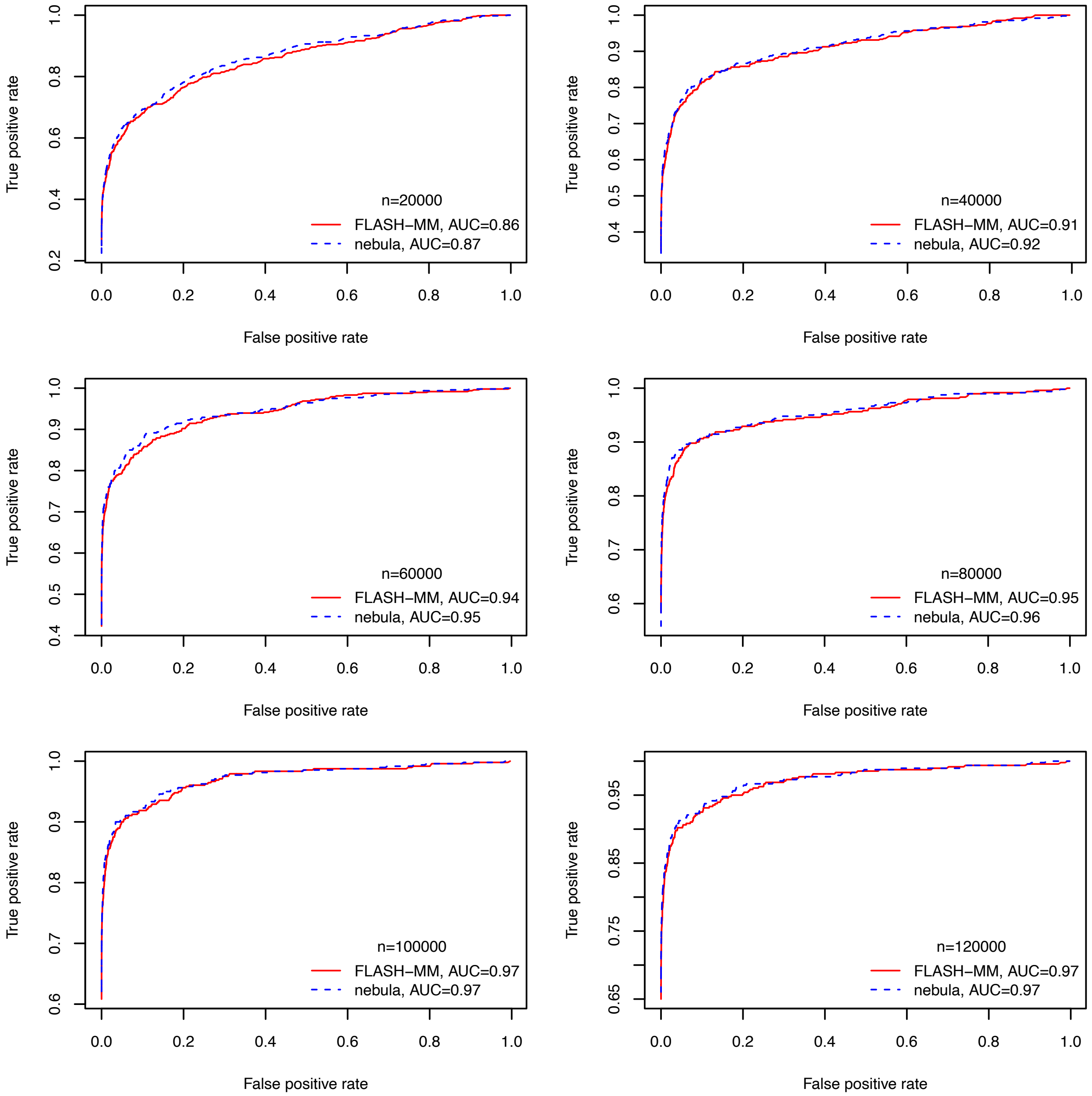


Extended Figure 2. ROC curves of TPR versus FPR for FLASH-MM and nebula in different sample sizes (n).

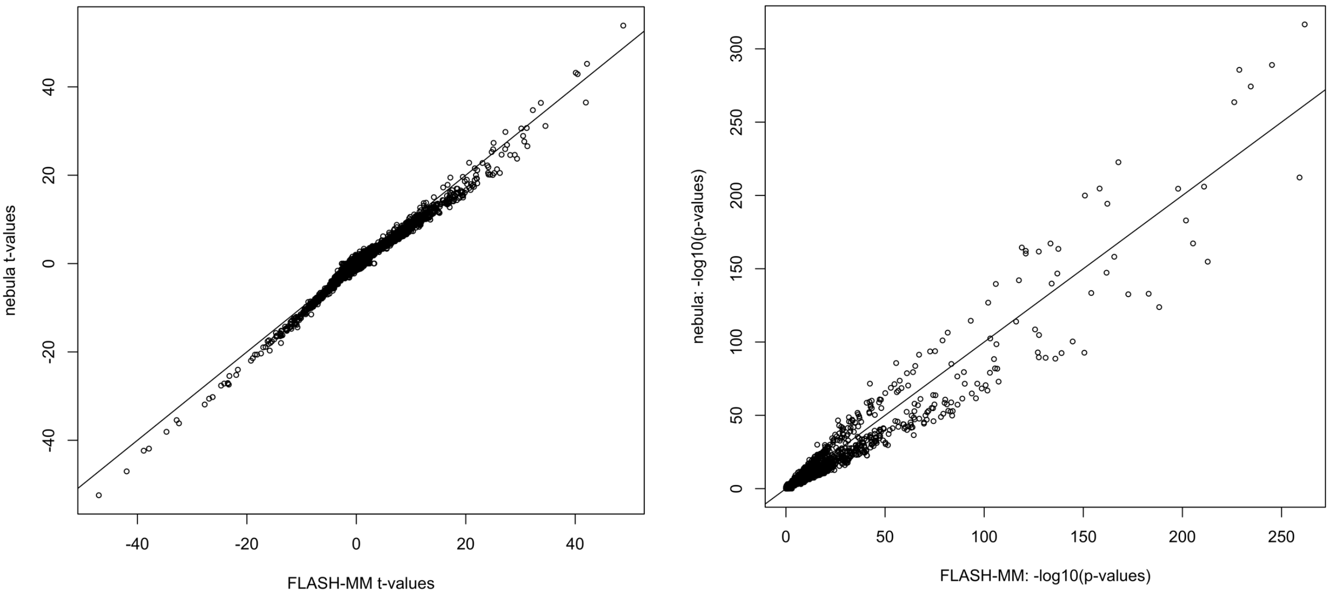


(A) t-values (B) p-values

Extended Figure 3. Scatterplots of FLASH-MM and nebula t-values and p-values.

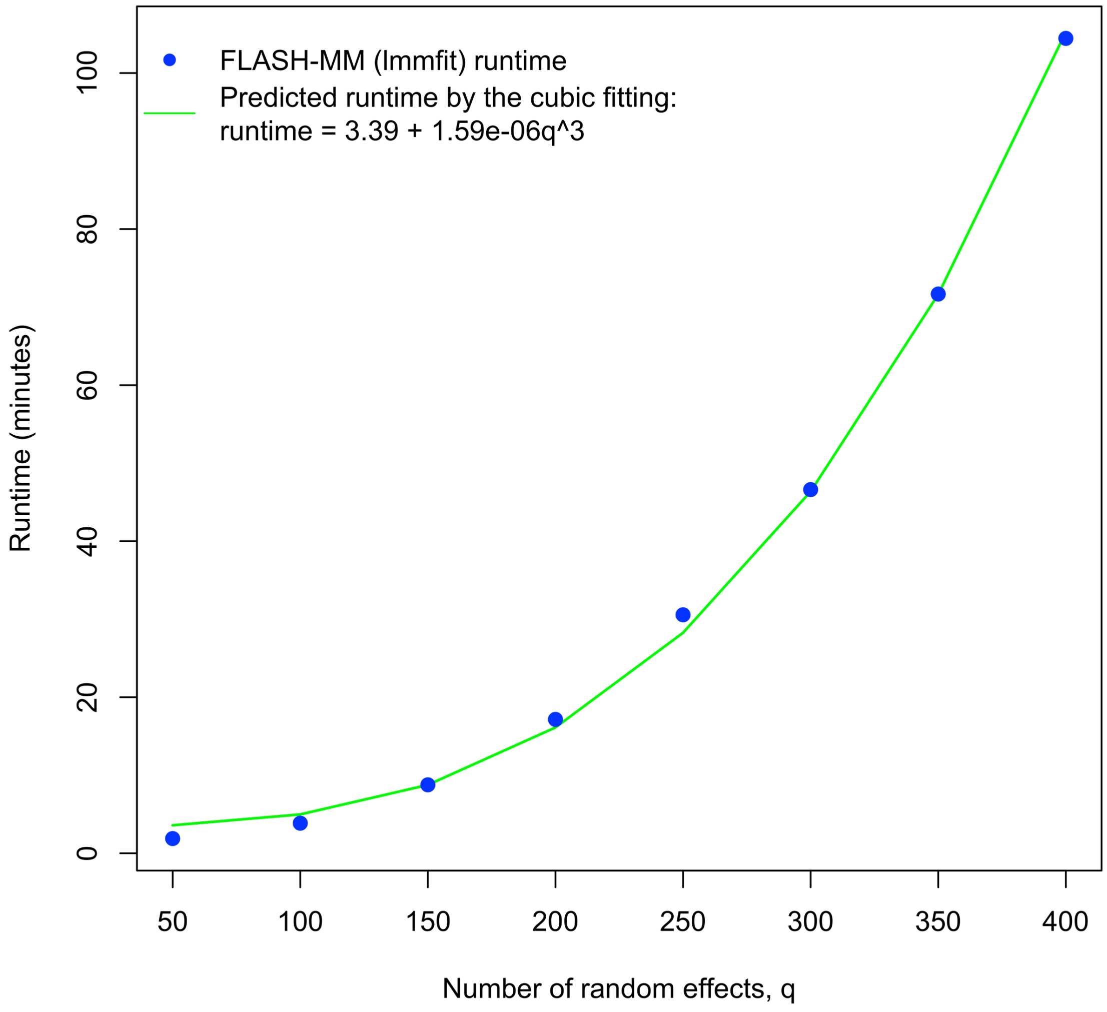


Extended Figure 4. Runtimes of the lmmfit function in the FLASHMM package for fitting the datasets containing 100,000 cells and 6,000 genes with 16 cell-types and 8 different numbers of individuals (random effects). The green curve is the predicted runtime by the cubic fitting function.
